# Mitf-family transcription factor function is required within cranial neural crest cells to promote choroid fissure closure

**DOI:** 10.1101/869040

**Authors:** Katie L. Sinagoga, Alessandra M. Larimer-Picciani, Stephanie M. George, Samantha A. Spencer, James A. Lister, Jeffrey M. Gross

**Author notes:** Corresponding Author: Jeffrey M. Gross.

## Abstract

A critical step in eye development is closure of the choroid fissure (CF), a transient structure in the ventral optic cup through which vasculature enters the eye and ganglion cell axons exit. While many factors have been identified that function during CF closure, the molecular and cellular mechanisms mediating this process remain poorly understood. Failure of CF closure results in colobomas. Recently, *MITF* was shown to be mutated in a subset of human coloboma patients, but how MITF functions during CF closure is unknown. To address this question, zebrafish with mutations in *mitfa* and *tfec*, two members of the Mitf-family of transcription factors, were analyzed and their functions during CF closure determined. *mitfa;tfec* mutants possess severe colobomas and our data demonstrate that Mitf activity is required within cranial neural crest cells (cNCCs) to facilitate CF closure. In the absence of Mitf function, cNCC migration and localization in the optic cup are perturbed. These data shed light on the cellular mechanisms underlying colobomas in patients with *MITF* mutations and identify a novel role for Mitf function in cNCCs during CF closure.

**Summary Statement:** Mitf-family transcription factors act within cranial neural crest cells to promote choroid fissure closure. Without Mitf-family function, cNCC localization and function in the CF is disrupted, thus contributing to colobomas.

## Introduction

Development of the vertebrate eye is a highly complex process, involving coordinated morphogenesis, migration, and communication between the surface ectoderm, neural ectoderm, and extraocular mesenchyme. During eye development, the optic primordia evaginate from the ventral forebrain and give rise to the optic vesicles (Bibliowicz et al., 2011). Subsequent invagination of the vesicle will give rise to the optic cup, which eventually subdivides into the retina and retinal pigmented epithelium (RPE). During optic cup morphogenesis the primitive optic stalk narrows ventrally and creates a transient opening, known as the choroid fissure (CF) (FitzPatrick, 2005; Schmitt and Dowling, 1994). The CF is critical for the entrance of the hyaloid vasculature into the eye and for exit of retinal ganglion cell axons. The CF must eventually close, such that retinal and RPE tissues are contained within the optic cup.

The CF is a remarkable area within the eye, in that it’s the point of convergence between the retina, RPE, and subtypes of periocular mesenchyme (POM). A network of signaling and regulatory pathways that includes Sonic Hedgehog and Retinoic Acid, among others, influences CF closure (Hyatt et al., 1996; Lupo et al., 2011; Marsh-Armstrong et al., 1994; Schimmenti et al., 2003; Take-uchi et al., 2003). The intersection of these major signaling pathways mediates transcription of downstream target genes within the CF, but also in the surrounding POM, which subsequently influence optic cup morphogenesis. The POM plays a crucial role in this process, as mesenchymal cells physically interact with and signal to the CF and optic cup to facilitate optic cup morphogenesis and CF closure (Bryan et al., 2018; Fuhrmann et al., 2000; Gestri et al., 2018; James et al., 2016). A large portion of POM cells are derived from the cranial neural crest (cNCC) (Williams and Bohnsack, 2015). Neural crest cells are born from the dorsal portion of the neural tube, after which they migrate ventrally and differentiate into the mesenchymal components of the face, including cartilage and vasculature (Bohnsack et al., 2011; Christiansen et al., 2000; Kaucka et al., 2016; Williams and Bohnsack, 2015). cNCCs additionally migrate anteriorly around the optic vesicle, during which they communicate with the optic stalk and retina (Grocott et al., 2011). Ultimately, morphogenetic and signaling events between the RPE, retina, and cNCC-derived-POM facilitate breakdown of basement membrane components within the fissure, such as laminin (Bryan et al., 2018; James et al., 2016; Lee and Gross, 2007), which enables fusion between the retina and RPE components of the CF. Despite these studies, the cellular and molecular underpinnings of CF closure remain unclear, with ECM interactions, signaling pathways, and a variety of transcription factors implicated in the process. Confounding this further, closure involves three distinct cell types – retina, RPE and POM.

When CF closure is perturbed, colobomas result. Colobomas are estimated to occur in 1 out of 10,000 live births and severe cases account for up to 10% of childhood blindness (Bermejo and Martínez-Frías, 1998; Onwochei et al., 2000; Stoll et al., 1997). Depending on where along the proximal-distal axis of the CF closure failed, one or several parts of the eye, including the lens, cornea, retina, optic nerve, and ciliary body can be involved (Bernstein et al., 2018; Gregory-Evans et al., 2004). While some mutated genes have been implicated in the pathology of the disease, these loci only make up a small portion (∼20%) of reported cases (Chang et al., 2006; FitzPatrick, 2005; Gregory-Evans et al., 2004). Recently, mutations were identified in *MITF*, a member of the microphthalmia-associated transcription factor/TFE (MiT) family of transcription factors (TFs), which result in colobomas (George et al., 2016). Patients with compound heterozygous mutations in *MITF* display a phenotype termed COMMAD syndrome (<u>C</u>oloboma, <u>O</u>steoporosis, <u>M</u>icrophthalmia, <u>M</u>acrocephaly, <u>A</u>lbinism and <u>D</u>eafness). MITF TFs are well-known for their roles in pigmentation, melanocyte development and specification of the RPE (Bharti et al., 2012; Hsiao and Fisher, 2014; Lister, 1999; Martina et al., 2014). Dominant-negative mouse Mitf mutations (*Mitf*^*mi/mi*^) result in colobomatous microphthalmia (Hero, 1989; Takebayashi et al., 1996), and loss of MITF in human embryonic stem cells impairs proliferation in optic vesicles differentiated from these cells (Capowski et al., 2014). Biochemically, *MITF* mutations from COMMAD patients result in decreased localization of MITF to the nucleus and decreased DNA binding ability to M-box and E-box target sites (George et al., 2016). However, despite these biochemical results, virtually nothing is known about the downstream mechanisms of how MITF contributes to CF closure or how *MITF* mutations result in colobomas.

The MITF family of transcription factors includes four members: Mitf, Tfec, Tfeb, and Tfe3 (Martina et al., 2014; Steingrímsson et al., 2004; Zhao et al., 1993). Members of the Mitf family homodimerize or heterodimerize with one another and bind to E-box and M-box DNA regions to control transcription of downstream targets (Hemesath et al., 1994; Pogenberg et al., 2012). Mitf is the most well studied member of the family, with key functions identified in cell cycle regulation, motility, metabolism, cell survival, and pigmentation (Cavodeassi and Bovolenta, 2014; Hsiao and Fisher, 2014). In addition, to paralogous genes in zebrafish (*mitfa* and *mitfb*), there are at least nine isoforms of Mitf/mitf, each maintaining a specific expression pattern during development in mouse and zebrafish (Bharti et al., 2008; Lister et al., 2001). For example, in mouse, Mitf-A regulates pigmentation in hair and tyrosinase expression in the eye, while Mitf-M functions within the kidney (Flesher et al., 2019). Many mitf isoforms have overlapping expression in zebrafish, but *mitfa* is expressed specifically in cNCCs, neural crest-derived melanocytes and the developing RPE (Lister et al., 2001). The MiTF family member *tfec* has a similar expression pattern to *mitfa* in the eye (Lister et al., 2011). While expressed in the RPE and cNCCs during early eye development, *tfec* is also enriched in the CF during critical closure time points (Cao et al., 2018). Due to the convergence of RPE and cNCC-derived POM in this region, it remains unknown in which cell type(s) Mitf transcription factors function to promote CF closure.

Ocular development and morphogenesis is highly conserved between zebrafish and humans, making this an ideal system to study CF closure. Utilizing a novel *mitfa;tfec* mutant line, we first demonstrate that loss of these Mitf-family transcription factors phenocopies colobomas observed in *MITF* patients. Our data indicate that both RPE and cNCC development is perturbed in *mitfa;tfec* mutants. Through a series of embryological manipulations and rescue experiments, we demonstrate that Mitf-family function is required specifically within the cNCCs to facilitate CF closure. Further, our data indicate that Mitf transcription factors act within cNCCs to promote migration into the eye and cell survival. Taken together, these data identify potential cellular underpinnings of colobomas in human COMMAD patients with mutations in *MITF* and provide a platform through which cNCC-specific functions during CF closure can be further elucidated.

## Results and Discussion

Previous studies have shown that *mitfa*^*-/-*^ mutants possess pigmentation defects, but ocular development is normal (Lister, 1999). Mitf-family members are co-expressed in many tissues and in zebrafish, *mitfa* and *tfec* have similar expression patterns within the RPE and cNCCs (Lister et al., 2001; Lister et al., 2011). Therefore, using CRISPR/Cas9, we generated a *tfec* mutant for analysis (Figure S1). The *vc60* allele contains a frameshifting indel in exon 7, encoding the second helix of the dimerization domain, resulting in a complete loss of function (Petratou et al., 2019). *tfec*^*-/-*^ mutants do not live beyond larval stages (1-2 weeks), as their swim bladders fail to inflate. *tfec*^*-/-*^ mutants possess pigmentation defects and are mildly microphthalmic, but overall optic cup formation is normal and they display no signs of colobomas (Fig. S1).

Considering that there may be some developmental compensation and/or functional redundancy between these genes, we next generated *mitfa*^*-/-*^*;tfec*^*-/-*^ double mutants and assessed eye development. At 4 days post fertilization (dpf), *mitfa*^*-/-*^*;tfec*^*-/-*^ mutants display prominent bilateral colobomas, that in many cases resulted in retinal “blowout” where retina and RPE were extruded into the forebrain (Fig. 1A). Only *mitfa*^*-/-*^*;tfec*^*-/-*^ mutants possessed colobomas; CF closure was normal in *mitfa*^*-/-*^*;tfec*^*+/+*^ (Fig. 1A) and *mitfa*^*-/-*^*;tfec*^*+/-*^ (data not shown) embryos. *mitfa*^*-/-*^*;tfec*^*+/-*^ mutants were viable and therefore for the remainder of the experiments, *mitfa*^*-/-*^*;tfec*^*+/-*^ incrosses were utilized to generate *mitfa*^*-/-*^*;tfec*^*-/-*^ mutants for analyses. Through this breeding scheme, expected Mendelian ratios of mutant embryos were recovered and colobomas appeared on average in 21% of progeny (Fig. S2). Colobomas vary in their severity in most models of the disease, as well as in human patients, and we likewise observed a range of severities in *mitfa*^*-/-*^*;tfec*^*-/-*^ mutants (Fig. 1B).

**Figure 1.**
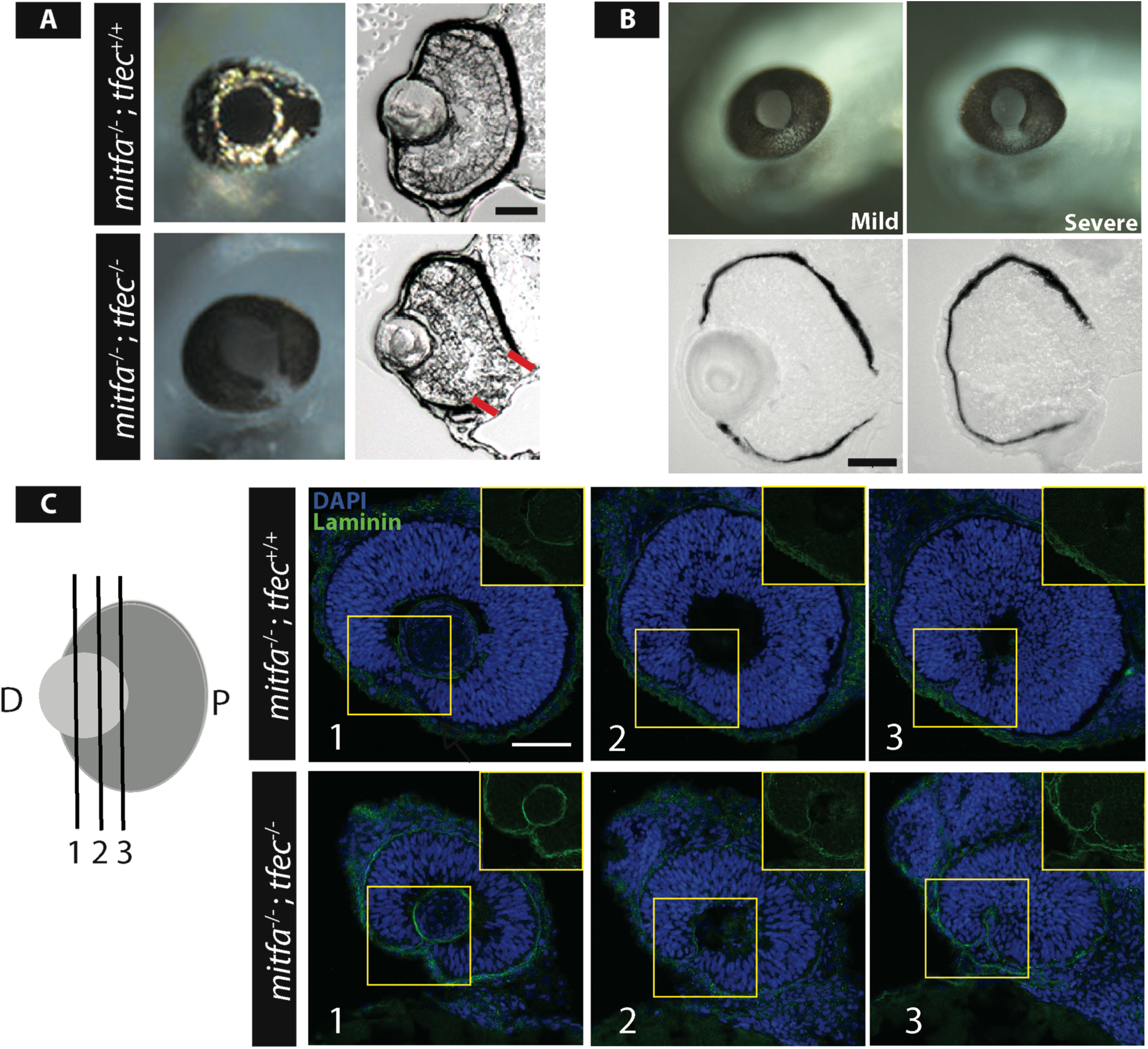
Zebrafish *mitfa*;*tfec* mutants phenocopy human colobomas. **A)** At 4dpf (days post fertilization), *mitfa*^*-/-*^*;tfec*^*-/-*^ mutants possess colobomas (outlined in red). **B)** At 4dpf, *mitfa*^*-/-*^*;tfec*^*-/-*^ mutants display a range of coloboma severities. **C)** At 48hpf, serial sections through the distal to proximal axis of the CF reveal persistent expression of laminin in the CF of *mitfa*^*-/-*^*;tfec*^*-/-*^ mutants. Scale bar = 50μm.

CF closure is a multistep process that relies on breakdown of the basement membrane (BM) lining the opposing sides of the CF prior to fusion (Bernstein et al., 2018; Carrara et al., 2019; James et al., 2016). A hallmark of colobomas in many models of the disease is BM retention within the CF (Barbieri et al., 2002; Geeraets, 1976; Hero, 1990; Hero et al., 1991; James et al., 2016; See and Clagett-Dame, 2009; Tsuji et al., 2012). Using laminin as a marker of the BM, we assayed BM degradation in *mitfa*^*-/-*^*;tfec*^*-/-*^ mutants. *mitfa*^*-/-*^*;tfec*^*-/-*^ mutants retain strong expression of laminin at the opposing sides of the CF, while *mitfa*^*-/-*^*;tfec*^*+/+*^ controls have degraded the intervening BM by 48hpf (Fig. 1C).

*mitfa* and *tfec* are expressed in two distinct cellular populations within the developing eye: the RPE and cNCCs (Lister et al., 1999; Lister et al., 2011). Activity in one or both of these tissues could mediate functions during CF closure that are disrupted in COMMAD patients. With this in mind, we examined the development of these two cell types in *mitfa*^*-/-*^*;tfec*^*-/-*^ mutants. At 26hpf, there was a dose-dependency to RPE pigmentation; the RPE of “control” *mitfa*^*-/-*^*;tfec*^*+/+*^ embryos was normally pigmented, while in *mitfa*^*-/-*^*;tfec*^*+/-*^ mutants it was hypopigmented and in *mitfa*^*-/-*^*;tfec*^*-/-*^ mutants it lacked nearly all pigmentation (Fig. 2A). However, these differences in RPE pigmentation resolved by 4dpf (data not shown). To examine cNCC development in *mitfa*^*-/-*^*;tfec*^*-/-*^ mutants, we utilized a *mitfa*:GFP transgene that labels a subset of migratory neural crest cells but not RPE (Curran et al., 2009). At 24hpf, in control embryos there were 0.6813 +/-0.051 neural crest cells per 1000μm^2^ area of the optic cup, whereas mutants possessed 0.3658 +/-0.029 neural crest cells per 1000μm^2^ area of the optic cup (p=0.0007, Fig. 2B). Of note, when we examined sections posterior to the eye, *mitfa*:GFP^+^ cNCCs were present near the neural tube (Fig. 2B). Transverse serial sections taken from *mitfa*^*-/-*^*;tfec*^*+/+*^*;mitfa*:GFP (control) and *mitfa*^*-/-*^*;tfec*^*-/-*^*;mitfa*:GFP mutants show substantially fewer cNCCs within the POM of *mitfa*^*-/-*^*;tfec*^*-/-*^ mutants at 48hpf as well (Fig. 2C). To address the possibility of developmental delay, whole-mount imaging of *mitfa*^*-/-*^*;tfec*^*+/-*^*;mitfa*:GFP (control) and *mitfa*^*-/-*^*;tfec*^*-/-*^*;mitfa*:GFP mutants was performed (Fig. S3A). Embryos lacked cNCCs surrounding the optic cup at 48 and 72hpf, suggesting that cNCC contribution to the POM is not simply delayed in *mitfa*^*-/-*^*;tfec*^*-/-*^ mutants.

**Figure 2.**
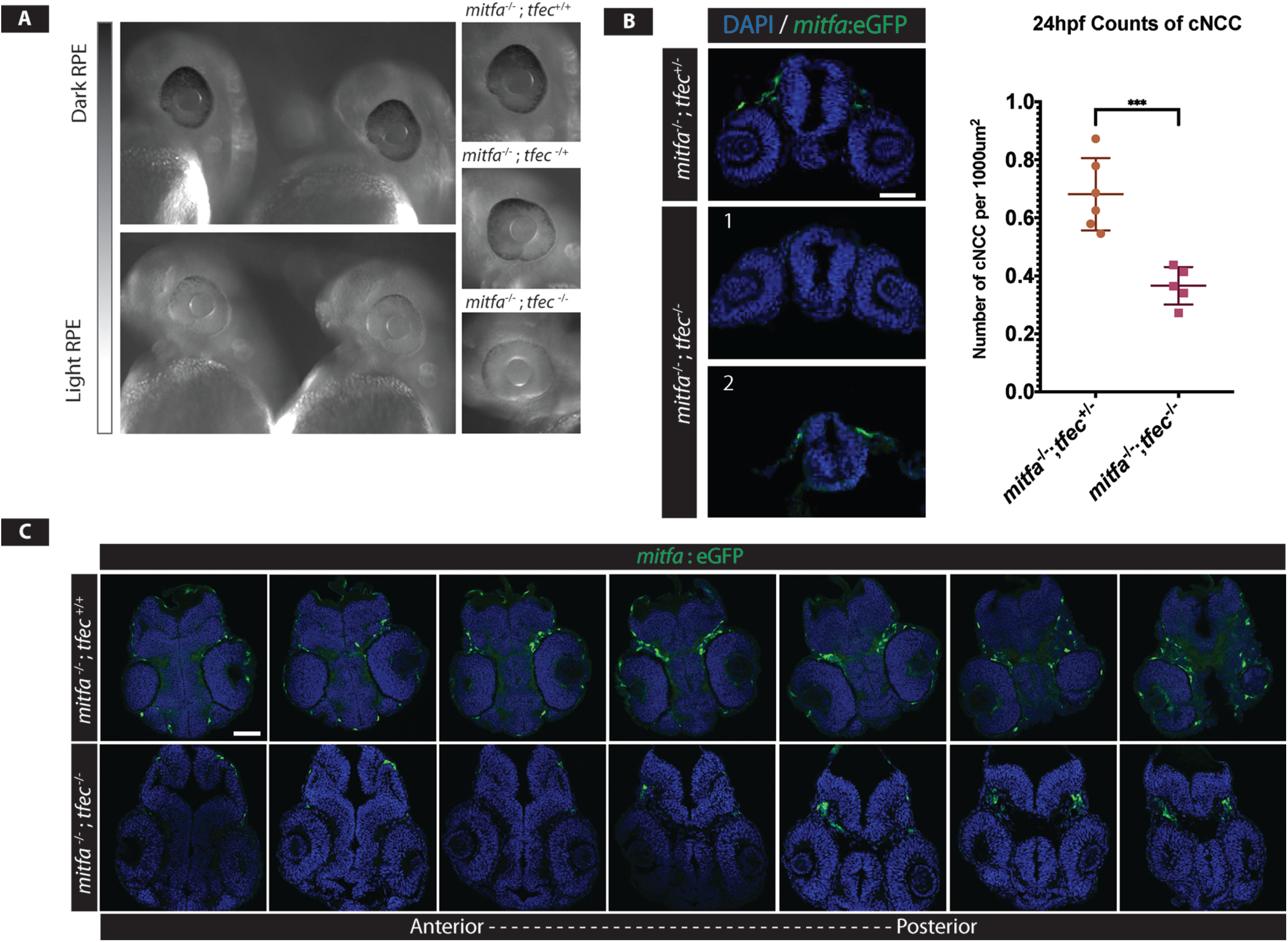
*mitfa* ^*-/-*^ *tfec* ^*-/-*^ mutants display RPE and cNCC phenotypes. **A)** At 26hpf, *mitf*^*-/-*^*;tfec*^*+-/-*^ and *mitfa*^*-/-*^*;tfec*^*-/-*^ mutants display mild to severe hypopigmentation, respectively. The RPE eventually becomes normally pigmented by 4dpf in both genotypes. **B)** At 24hpf fewer cNCCs are present around the eye of *mitfa* ^*-/-*^; *tfec* ^*-/*^; *mitfa*:GFP embryos (p=0.0007). *mitfa* ^*-/-*^*;tfec* ^*-/-*^ mutants contain cNCCs posterior to the optic field, however (2). **C)** Serial sections of *mitfa* ^*-/-*^*;tfec* ^*+/+*^;*mitfa:*GFP and *mitfa* ^*-/-*^*;tfec* ^*-/-*^;*mitfa:*GFP embryos demonstrate that *mitfa* ^*-/-*^*;tfec* ^*-/-*^ mutants possess fewer cNCCs in the POM surrounding the eyes. Scale bar = 50μm.

Optic cup invagination and CF formation occurs from 16-24hpf (Schmitt and Dowling, 1994). At the same time, cNCCs are being specified from the dorsal neural tube and will migrate anteriorly around the eye (Christiansen et al., 2000), with the eye thought to provide a source of attractant for their migration (Langenberg et al., 2008). Given the decrease in cNCC numbers in the POM of *mitfa*^*-/-*^*;tfec*^*-/-*^ mutants, we predicted that migration of cNCCs was perturbed in *mitfa;tfec* mutants. Utilizing *in vivo* 4D-time-lapse confocal microscopy, we followed the migration of *mitfa:*GFP expressing cNCCs in wild type and mutant embryos for 15 hours starting at 25hpf (Supp. Videos 1,2). *mitfa*^*-/-*^*;tfec*^*-/-*^ mutant cNCCs did not migrate from the dorsal neural tube to the eye. From the time lapse data, some cNCCs in *mitfa*^*-/-*^*;tfec*^*-/-*^ mutants appeared to explode (see t=8:08-10:08 in Supp Video 2), consistent with cell death during their migration. To directly assess whether elevated apoptosis contributed to the lack of cNCCs around the eye of *mitfa*^*-/-*^*;tfec*^*-/-*^ mutants, we performed TUNEL staining at 24hpf. While the number of TUNEL^+^ cNCCs was elevated in *mitfa*^*-/-*^*;tfec*^*-/-*^ mutants, the difference was not significant when compared to wild-type controls, suggesting that lack of cNCCs within the POM in *mitfa*^*-/-*^*;tfec*^*-/-*^ mutants is primarily driven by a defect in cell migration (Fig. S3B). Combined, these data demonstrate a transient delay in RPE pigmentation in the absence of mitfa/tfec function and a significant defect in cNCC localization to the POM in *mitfa*^*-/-*^*;tfec*^*-/-*^ mutants.

Ocular hypopigmentation or albinism is a common feature of a number of human disorders and these do not commonly correlate with colobomas, nor do most defects in RPE formation/function result in colobomas (e.g. Ma et al., 2019; Fuhrmann 2010; Reissman and Ludwig, 2013; Jeffrey 1998). On the other hand, there are many studies demonstrating functional requirements for cNCCs and POM in regulating early eye development and distinct aspects of CF closure (Akula et al., 2019; Bryan et al., 2018; Dee et al., 2013; Fuhrmann et al., 2000; Gestri et al., 2018; James et al., 2016; Lupo et al., 2011; McMahon et al., 2009; Sedykh et al., 2017). While our imaging data demonstrate abnormalities in the cNCC population of *mitfa*^*-/-*^*;tfec*^*-/-*^ mutants, given that Mitf-family genes are expressed in both RPE and cNCCs, it remains formally possible that defects within the RPE could also contribute to colobomas. Thus, we wanted to directly assess in which population *mitfa* and *tfec* function was required to close the CF. Zebrafish are highly amenable to embryonic transplantation experiments, in which cells can be transplanted between embryos early in development to create genetic mosaics (Carmany-Rampey and Moens, 2006), with cells targeted to specific tissues or cell types based on established fate maps (Woo and Fraser, 1995), to determine in which cell type(s) a particular gene product functions.

To test the hypothesis that Mitf-family function was required in cNCCs during CF closure, we transplanted wild-type cells into either the cNCC domain (Fig. 3A) or the retina/RPE domain (Fig. 3C) of *mitfa*^*-/-*^*;tfec*^*-/-*^ mutant host embryos at shield stage and quantified CF closure at 4dpf. Donor embryos were injected with fluorescent dextran at the 1 or 2-cell stage to track their progeny in hosts and validate correct targeting. Wild-type cells transplanted into the domain that gives rise to cNCCs in *mitfa*^*-/-*^*;tfec*^*-/-*^ mutants resulted in a population of strongly labeled cNCCs at 24hpf (Fig. 3A). Remarkably, wild-type cNCCs rescued CF closure in *mitfa*^*-/-*^*;tfec*^*-/-*^ mutants (Fig. 3B). Qualitatively, the CF appeared fully closed in 6 of 7 transplanted mutants when examined intact (Fig. 3B), while quantitatively, the maximum angle of CF opening in serial sections from non-transplanted mutant eyes was 26.9 +/-3.04 degrees, while in *mitfa*^*-/-*^*;tfec*^*-/-*^ mutants containing wild-type cNCCs, it was reduced to 10.33 +/-2.65 degrees (p=0.0004; Fig. 3E). Transplantation of wild-type cells into the cNCC domain of *mitfa*^*-/-*^*;tfec*^*+/-*^ embryos had no effect (Fig. S4A). These data indicate that Mitf-family function in cNCCs is sufficient to mediate CF closure in in *mitfa*^*-/-*^*;tfec*^*-/-*^ mutants. Wild-type cNCCs also retained the ability to migrate to the eye of *mitfa*^*-/-*^*;tfec*^*-/-*^ mutants (Fig. 3B), supporting a cell autonomous role for *mitfa* and *tfec* within cNCCs in mediating their migration to the eye.

**Figure 3.**
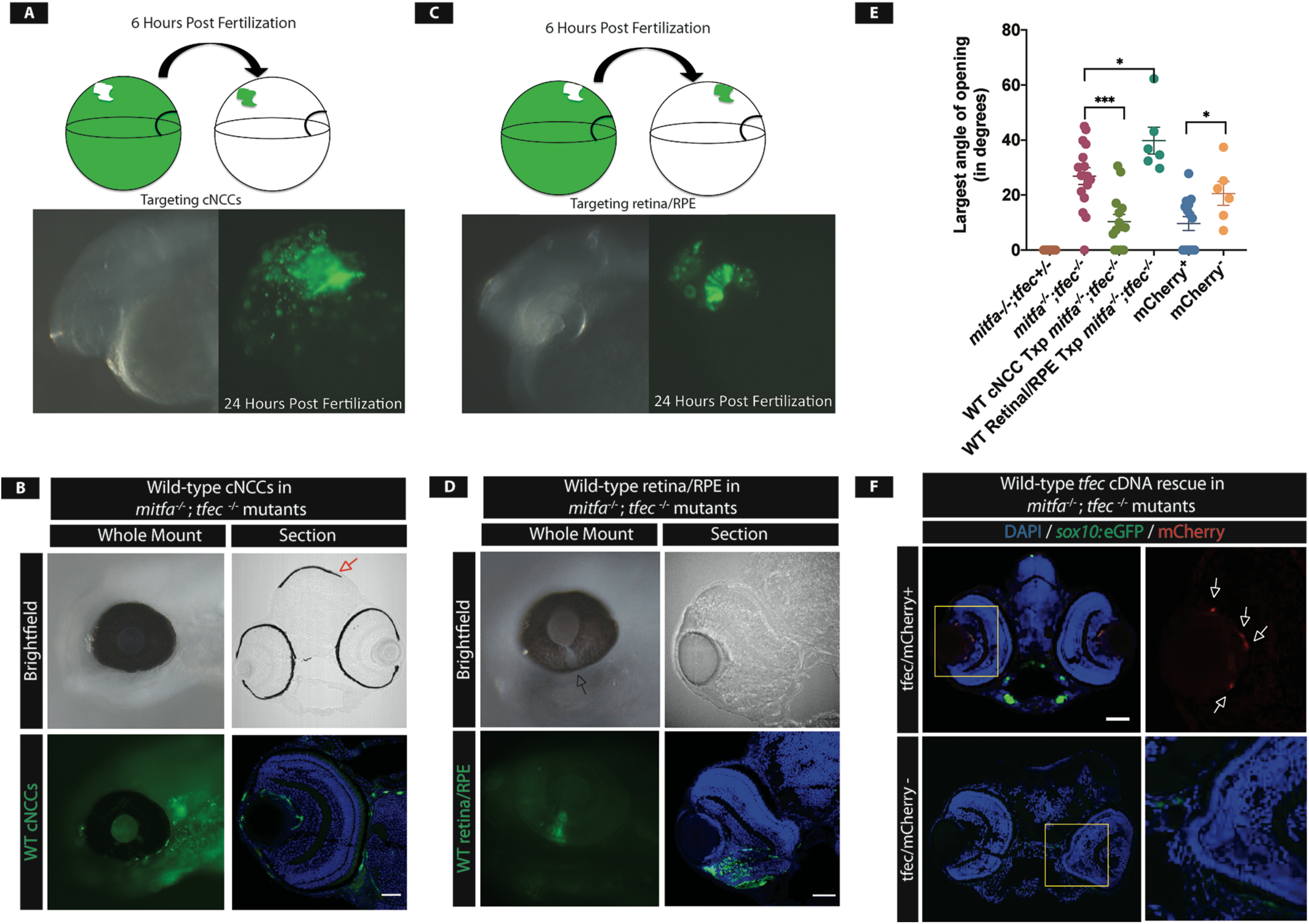
*mitfa* and *tfec* are required within cNCCs to promote CF closure. **A)** Neural crest transplantations. At 6hpf, 10-15 wild-type, dextran labeled cells are transplanted ∼90 degrees off the organizer in a *mitfa*^*-/-*^*;tfec*^*-/-*^ embryo. At 24hpf, transplanted cells, marked by FITC-dextran, are detected in the cNCC population, indicating successful transplantation. **B)** Examples of 4dpf wild-type cNCC transplanted *mitfa*^*-/-*^*;tfec*^*-/-*^ embryos where the coloboma has been rescued. Transplanted cNCCs are detected in and around the eye and rescue of a cNCC-derived pigmented melanocyte (red arrow) confirms successful transplantation (scale bar = 25μm) **C)** Retina/RPE transplants. At 6hpf, wild-type, FITC-dextran labeled cells are transplanted into the RPE/retinal region of a *mitfa*^*-/-;*^*tfec*^*-/-*^ mutant. At 24hpf, wild-type cells are detected within the developing optic cup, indicating successful transplantation. **D)** At 4dpf, transplanted embryos still possess colobomas. **E)** Quantification of colobomas (largest angle of CF opening) in rescue experiments. CF opening is significantly reduced in *mitfa*^*-/-;*^*tfec*^*-/-*^ embryos transplanted with wild-type cNCCs (p=0.0004, n=14 eyes). However, colobomas are not resolved in retina/RPE transplanted mutants and are actually larger than in non-transplanted control *mitfa*^*-/-;*^*tfec*^*-/-*^ mutants (p=0.037, n=6 eyes). *tfec* expression (mCherry^+^) also rescues colobomas (p=0.033, n=14 eyes) **F)** pDestTol2pA2-*sox10*:mCherry-nostop-t2A-FLAG-*tfec*-pA injection reduces coloboma size in *mitfa*^*-/-;*^*tfec*^*-/-*^ mutants. Scale bar = 50μm.

We next tested the alternative hypothesis, that Mitf-family function was required in the retina/RPE to enable CF closure. Wild-type cells transplanted into the domain that gives rise to retina/RPE in *mitfa*^*-/-*^*;tfec*^*-/-*^ mutants resulted in a strongly labeled optic cup at 24hpf, and only embryos in which at least 20% of the optic cup was composed of transplanted cells were utilized for subsequent analyses (Fig. 3C). Despite robust transplantation, wild-type retinal/RPE cells failed to rescue colobomas in *mitfa*^*-/-*^*;tfec*^*-/-*^ mutants, even in cases where they were largely restricted to the ventral optic cup (Fig. 3D). The maximum angle of CF opening in sections from non-transplanted mutant eyes was 26.9 +/-3.04 degrees, while in *mitfa*^*-/-*^*;tfec*^*-/-*^ mutants containing wild-type retina/RPE, it was 39.82 +/-4.87 degrees (p=0.037; Fig. 3E). Transplantation of wild-type cells into the retina/RPE of *mitfa*^*-/-*^*;tfec*^*+/+*^ embryos had no effect (Fig. S4B). These data indicate that Mitf-family function in the retina/RPE does not likely facilitate CF closure. A caveat here is that our transplants could be sub-threshold for the number of cells necessary to mediate effective closure. However, as rescue did not occur even in cases where wild-type cells were largely restricted to the ventral optic cup and CF region (Fig. 3D), our data support a model in which Mitf-family function in cNCCs mediates CF closure.

To further test this model, we expressed wild-type *tfec* specifically within cNCCs of *mitfa*^*-/-*^*;tfec*^*-/-*^ mutants and assessed CF closure. *tfec* is enriched at the CF during CF closure (Cao et al., 2018) and wild-type *tfec* is sufficient for CF closure in *mitfa*^*-/-*^ mutants (Figs. 1 and S1). Utilizing the Tol2 system (Kwan et al., 2007) we generated pDestTol2pA2-*sox10*:mCherry-nostop-t2A-FLAG-*tfec*-pA and injected embryos derived from *mitfa*^*-/-*^*;tfec*^*+/-*^*;sox10:*eGFP incrosses. Embryos that possessed detectable mCherry^+^ cNCCs were assessed at 4dpf for colobomas, sectioned serially and the maximum angle of CF opening quantified. Expression of wild-type *tfec* in *mitfa*^*-/-*^*;tfec*^*-/-*^ mutants reduced the CF opening from 20.58 +/-4.31 to 9.66 +/-2.49 (p=0.033; Fig. 3E,F).

In summary, static and *in vivo* imaging data identified defects in cNCC migration in *mitfa*^*-/-*^*;tfec*^*-/-*^ mutants and defects in breakdown of the BM lining the CF, functions attributed to the POM during CF closure (Bryan et al., 2018; Fuhrmann et al., 2000; Gestri et al., 2018; James et al., 2016). Data from cell transplantation and cDNA rescue experiments demonstrate that Mitf-function in cNCCs is sufficient to rescue CF closure defects in *mitfa*^*-/-*^*;tfec*^*-/-*^ mutants. Taken together, these data strongly support a model in which Mitf-family function is required in cNCCs to facilitate CF closure. Without sufficient cNCC contribution/function within the POM, colobomas result. These data are consistent with a model arising from other studies implicating cNCCs as critical regulators of optic cup morphogenesis and early eye development (Bryan et al., 2018; Fuhrmann et al., 2000). Additionally, they highlight the expanding role of the MiTf family of transcription factors in a variety of developmental processes beyond their well-known roles in regulating pigmentation (Hodgkinson et al., 1993; Price and Fisher, 2001; Steingrímsson et al., 1994). Finally, these data identify a potential cellular mechanism underlying colobomas in COMMAD patients with *MITF* mutations (George et al., 2016). Indeed, several human congenital disorders/syndromes that include colobomas also have phenotypes consistent with neural crest defects (e.g. Akula et al., 2019; Asad et al., 2016) and further studies can now focus on the neural crest origin of ocular diseases and how cNCCs modulate optic cup morphogenesis and CF closure. Future studies addressing these developmental questions will be essential to understanding the link between neural crest functions and congenital disorders of eye formation.

## Materials and Methods

### Zebrafish Husbandry

Zebrafish were maintained on a 14 hour/10 hour light-dark cycle at 28.5°C. For the *mitfa* and *tfec* lines, fish were maintained in a *mitfa*^-/-^ mutant background and double mutant embryos were obtained by mating pairwise heterozygous *tfec*^+/-^ crosses. The mutant alleles *mitfa*^*w2*^ (Lister et al. 1999) and *tfec*^*vc60*^ (Petratou et al. 2019) were used for all experiments. Wild type AB animals were used as donors for rescue transplantation experiments. Genotyping of fish was done using High Resolution Melt Analysis (HRM) using primers for tfec: Forward (GTGATATGCGCTGGAACAAAGGGA) Reverse (GCTCTTTCTGCAGCCACTTAATGTAT). *sox10*:eGFP fish were previously generated (Hoffman et al., 2007) and obtained from Dr. Ann Morris (University of Kentucky). All fish were housed and maintained in accordance with the University of Pittsburgh School of Medicine Institutional Animal Care and Use Committees.

### Tissue Preparation and Cryosectioning

Embryos were collected and fixed in 4% PFA in PBS for 1 hour at room temperature. Embryos were subsequently washed with 25% and 35% sucrose in PBS and embedded in Tissue Plus O.C.T Compound (Fisher Scientific). Tissue blocks were frozen and maintained at -80°C. Tissue was sectioned at 12μm on poly-lysine coated FrostPlus slides (Fisher). Slides were maintained at 4°C until analysis.

### Immunohistochemistry

Slides were rehydrated with 3× 20-minute washes of 1x PBS. For laminin staining, antigen retrieval was performed by incubation in 0.5% SDS at 37°C for 20 minutes. Slides were washed in PBS and then blocked using 10% normal goat serum in PBS-TD for 1 hour. Staining was performed overnight at 4°C in blocking solution. The antibodies used in this study were as follows: laminin*α*1 (Sigma-Aldrich) 1:100, DAPI (Life Technologies) 1:100, goat anti rabbit Alexa 488 secondary (Jackson) 1:500.

### 24hpf cNCC quantification and TUNEL Assay

*mitfa*:GFP embryos were collected at 24 hours post fertilization and fixed in 4% PFA in PBS for 1 hour at room temperature. Genotyping and tissue preparation of embryos was performed as outlined above. Tissue was sectioned at 12μm and In Situ Cell Death Detection Kit, TMR Red (Roche) was used to assess cell death. Tissue was also stained with DAPI (Life Technologies) 1:100. Cell death and quantification of total neural crest cells was completed counting the total number of cranial *mitfa*:GFP cells and *mitfa*:GFP^+^ TUNEL^+^ cells in each half of the embryo. Numbers were normalized to the area of the eye in the respective side of the head and results are displayed as the number of neural crest cells/per 1000μm^2^ area of the eye.

### Imaging

Slides were imaged using an Olympus FV1200 confocal microscope using Olympus software. Ten to twelve 1μm optical sections were acquired and then stacked using ImageJ software.

Whole mount imaging was performed using an Olympus FV1200 confocal microscope. Embryos were mounted in 0.5% low melt agarose and then immersed in Danieu’s Embryo Media. Embryos were lived imaged for 15 hours starting at 25hfp and analyzed using Olympus software.

### Transplantation Assays

Fish lines were bred as outlined above. Donor embryos were injected with 3.5% Alexa Fluor Dextran 488 (Thermo Fisher Scientific). Embryos were maintained in Danieus’s Embryo media and de-chorionated at 4hpf. At 6hpf, embryos were embedded in a 1.5% Methylcellulose and 2 drops of Ringer’s High Calcium. For transplant rescue experiments, 10-15 dextran-positive cells of a wild-type embryo were transferred into the region of neural crest or retinal/RPE origin in a *mitfa*^-/-^; *tfec*^-/-^ embryo. Embryos were then maintained in sterile filtered Danieu’s embryo media with 1x Pen/Strep for 24 hours. At 24 hours, embryos were assessed for successful transplantation using a Zeiss Axio Zoom V16 Dissecting Scope and Z3 Zeiss software. Embryos were grown until 4dpf, at which point they were assessed for the presence or absence of colobomas. Preparation of tissue included collection, fixation, transverse sectioning and imaging as described previously. Images were analyzed for the presence and quantification of CF opening. To quantify CF opening, ImageJ was used to define a line between the dorsal and ventral inner plexiform boundaries of the retina. From the central point on this line, the angle of opening of the CF was calculated from each serial section.

### cDNA injections

The plasmid pDestTol2pA2-*sox10*:mCherry-nostop-t2A-FLAG-*tfec*-pA used for cDNA microinjection was generated by multi-site Gateway cloning (Invitrogen/Thermo-Fisher Scientific) from the plasmids pDestTol2pA2 (Kwan et al. 2007; Tol2Kit #394), p5E-MCS-sox10-4.8 (Prendergast et al., 2012)(doi: 10.1242/dev.072439), pME-mCherry-no stop (Kwan et al. 2007; Tol2Kit #456), and p3E-2A-FLAG-tfec-pA (Petratou et a. 2019).

### Statistics

Student’s unpaired, two-tailed t-test was used to assess significance and error bars on all graphs indicate standard error of the mean (SEM).

## Acknowledgements

We thank Kira Lathrop and the University of Pittsburgh Ophthalmology Imaging Core for advice on imaging, Hugh Hammer of the University of Pittsburgh Fish Facility for help and guidance on fish maintenance and Corin Tucker, Carrie Brownstein and Janet Weiss for technical support.

## Figure Legends

**Figure S1. *tfec***^***-/-***^ **single mutants do not possess colobomas**. Whole mount and sections of 4dpf *tfec* ^*-/-*^ mutants reveals mild microphthalmia and pigmentation defects. No colobomas are detected. Scale bar = 100μm)

**Figure S2. Quantification of colobomas in *mitfa*** ^***-/-***^***;tfec***^***+/-***^ **incrosses. A)** Coloboma incidence in embryos at 4dpf from *mitfa* ^*-/-*^*;tfec* ^*+/-*^ incrosses. On average 21% of embryos possessed colobomas of varying severity. (n=4 rounds of breeding) **B)** Genotypes of embryos in *mitfa* ^*-/-*^*;tfec* ^*+/-*^ incrosses. Expected Mendelian ratios of *mitfa* ^*-/-*^*;tfec* ^*+/+*^, *mitfa* ^*-/-*^; *tfec* ^*+/-*^, and *mitfa* ^*-/-*^*;tfec* ^*-/-*^ are observed (n= 4 rounds of breeding).

**Figure S3. Loss of cNCCs in the POM is not the result cell death. A)** Whole-mount images of *mitfa* ^*-/-*^*;tfec* ^*+/-*^ and *mitfa* ^*-/-*^*;tfec* ^*-/-*^ embryos reveals persistent loss of cNCC within the POM at 48hpf and 72hpf. **B)** Quantification of TUNEL^+^ cNCCs. While a trend of increased cell death is present, loss of cNCC numbers cannot be attributed solely to apoptosis at 24hpf (p=0.512, n=5 embryos).

**Figure S4. Transplantation of wild-type cells does not affect eye development. A)** Wild-type cells transplanted into the region of cNCC origin in a *mitfa* ^*-/-*^*;tfec* ^*+/-*^ embryo shows no effect on eye development. **B)** Similarly, transplanted wild-types cells targeting the retina/RPE in *mitfa* ^*-/-*^*;tfec* ^*+/+*^ embryos show no effect on eye development.

**Supplementary Movies. Movie 1:** Wild-type embryos show normal migration of cNCCs starting at 25hpf. cNCCs can be seen migrating in and around the developing optic cup, eventually making their way around the entirety of the eye and into the CF. **Movie 2:** Mutant embryos show remarkably fewer cNCCs at 25hpf. Already, there is a lack of migration of the cells into and around the eye. These cNCCs never reach the CF during the 15-hour time-lapse movie and instead stay in the dorsal portion of the developing head. cNCCs seem to lack direction and at t=8:08-10:08, several dorsal cNCCs burst, implicating increased cNCC cell death in *mitfa* ^*-/-*^*;tfec* ^*-/-*^ embryos.

